# Inhibition of the angiotensin-converting enzyme N-terminal catalytic domain prevents endogenous opioid degradation in brain tissue

**DOI:** 10.1101/2025.01.21.634163

**Authors:** Filip Hanak, Jessica L. Swanson, Krzysztof Felczak, Prakashkumar Dobariya, Ursula C. H. Girdwood, Kenneth E. Bernstein, Swati S. More, Patrick E. Rothwell

**Author notes:** Corresponding author: Patrick E. Rothwell, 6-145 Jackson Hall, 321 Church St SE, Minneapolis, MN 55455.

## Abstract

Angiotensin-converting enzyme (ACE) regulates blood pressure by cleaving angiotensin peptides in the periphery, and can also regulate endogenous opioid signaling by degrading enkephalin peptides in the brain. ACE has two catalytic domains, located in the N-terminal or C-terminal region of the protein, but little is known about the roles that these two catalytic domains play in regulating endogenous opioid degradation in brain tissue. Using acute brain slice preparations from mice of both sexes, we developed methods to study degradation of Met-enkephalin-Arg-Phe (MERF) by ACE, and investigated the role of each ACE catalytic domain in MERF degradation. Using mutant mouse lines with functional inactivation of either the N-terminal domain or the C-terminal domain, we incubated acute brain slices with exogenous MERF at a saturating concentration. The degradation MERF to produce Metenkephalin was only reduced by N-terminal domain inactivation. Additionally, application of a selective N-terminal domain inhibitor (RXP407) reduced degradation of both exogenously applied and endogenously released MERF, while a selective C-terminal domain inhibitor (RXPA380) had no effect. Taken together, our results suggest that the ACE N-terminal domain is the primary site of MERF degradation in brain tissue, and that N-terminal domain inhibition is sufficient to reduce degradation of this specific endogenous opioid peptide. Our results have exciting implications for the development of novel pharmacotherapies that target the endogenous opioid system to treat psychiatric and neurological disorders.

---

Angiotensin-converting enzyme (ACE) is a dipeptidyl carboxypeptidase, cleaving two amino acids from the C-terminus of peptide substrates (1). ACE is well-known for regulating blood pressure by converting angiotensin I to angiotensin II. We recently identified another role for ACE in the brain (2): modulating synaptic plasticity by cleaving and degrading Met-enkephalin-Arg-Phe (MERF), an endogenous opioid peptide. ACE inhibition prevents MERF degradation (Supplemental Figure 1) and enhances endogenous opioid signaling in the nucleus accumbens, a brain region with conjointly high expression of ACE, MERF, and cognate opioid receptors. This could plausibly explain clinical reports that centrally active ACE inhibitors have unexpected secondary benefits (see Supplemental Text). However, the mechanism by which ACE cleaves MERF and regulates endogenous opioid signaling in brain tissue remains poorly understood.

**Figure 1.**
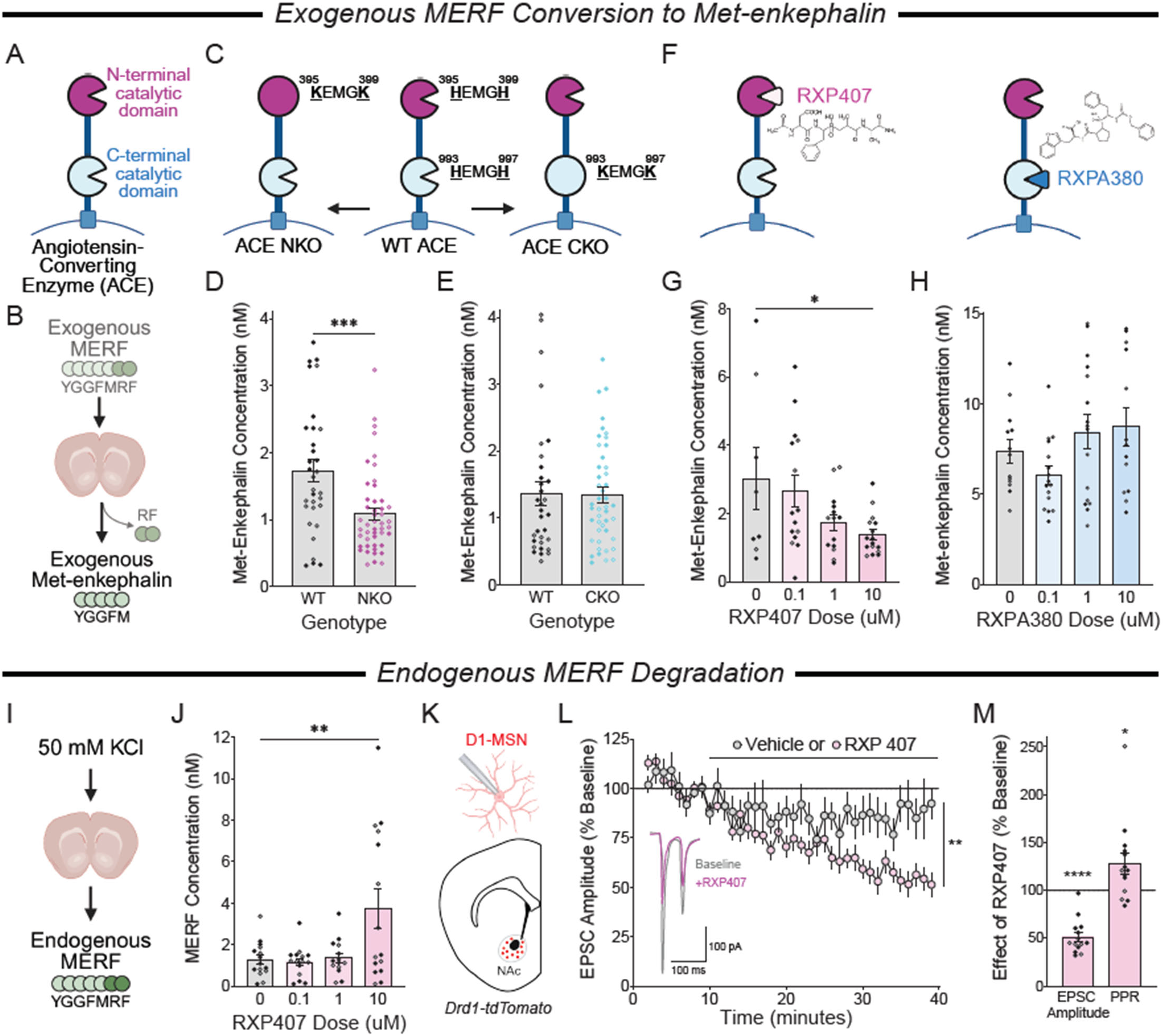
Met-enkephalin-Arg-Phe (MERF) degradation by the angiotensin-converting enzyme (ACE) N-terminal catalytic domain. (**A**) Illustration of ACE catalytic domains. (**B**) Experimental design: exogenous MERF applied to brain slices and cleaved by ACE to produce Met-enkephalin. (**C**) Genetic mutations inactivating the ACE N-terminal domain (“NKO”) or C-terminal domain (“CKO”). (**D**) MERF conversion to Met-enkephalin in NKO mice (n=47) and wild-type (WT) littermates (n=32). ANOVA Genotype main effect: F_1,75_=14.22, p=0.0003. (**E**) MERF conversion to Met-enkephalin in CKO mice (n=45) and WT littermates (n=32). ANOVA Genotype main effect: F_1,73_<1. (**F**) Pharmacological inhibition of ACE N-and C-terminal domains with RXP407 and RXPA380, respectively. (**G**) Effects of RXP407 on MERF conversion to Met-enkephalin (n=8-16/dose). ANOVA Dose main effect: F_3,45_=3.25; p=0.031; ^*^p<0.05, Dunnett’s post-hoc test. (**H**) Effects of RXPA380 on MERF conversion to Met-enkephalin (n=13-16/dose). ANOVA Dose main effect: F_3,54_=2.33, p=0.085. (**I**) Experimental design: stimulation of brain slices with 50mM KCl to release endogenous MERF. (**J**) Effects of RXP407 on MERF extracellular concentration (n=15-16/dose). ANOVA Dose main effect: F_3,51_=5.38; p=0.003; ^**^p<0.005, Dunnett’s post-hoc test. (**K**) Whole-cell patch-clamp recording from nucleus accumbens Drd1-expressing medium spiny neuron (D1-MSN). (**L**) Bath application of RXP407 (10µM, n=15) caused long-term depression of evoked excitatory postsynaptic current (EPSC) amplitude, which was significantly greater than vehicle (n=10). ^**^p<0.01, ANOVA Treatment x Time interaction. (**M**) Effects of RXP407 on EPSC amplitude and paired-pulse ratio. ^*^p<0.05, ^****^p<0.0001; one-sample t-test versus reference value (100%). Graphs display mean ± SEM, with individual data points from females and males shown as open and closed circles, respectively.

ACE has two catalytic domains, located in the N-terminal or C-terminal region of the protein (Figure 1A), with distinct profiles of substrate specificity (1). To investigate the contribution of each catalytic domain to MERF degradation, we used liquid chromatography-tandem mass spectrometry (LC-MS/MS) to quantify extracellular enkephalin in mouse brain tissue (2, 3). Acute coronal brain slices containing the nucleus accumbens were incubated in artificial cerebrospinal fluid containing a saturating concentration of exogenous MERF, which is cleaved and degraded by ACE to produce Met-enkephalin (Figure 1B). To study the role of each catalytic domain in MERF degradation, we used mouse lines carrying mutations in the active site of either the N-terminal catalytic domain (NKO) or the C-terminal catalytic domain (CKO; Figure 1C). Crucially, these mutations are amino acid substitutions that do not change the expression level of ACE and preserve the function of the intact domain (4, 5). The NKO mutation significantly reduced conversion of exogenous MERF to Met-enkephalin (Figure 1D), whereas the CKO mutation had no effect (Figure 1E). We complemented this genetic analysis with acute pharmacological inhibition using RXP407 and RXPA380, small molecule ACE inhibitors with a high selectivity for the N-terminal and C-terminal catalytic domains, respectively (Figure 1F). RXP407 reduced Met-enkephalin production in a dose-dependent manner (Figure 1G), recapitulating our findings with NKO mice, whereas RXPA380 had no effect (Figure 1H).

To build on our analysis of exogenous MERF degradation by ACE, we next measured the degradation of endogenous MERF released from brain tissue following chemical stimulation with a high concentration of potassium chloride (Figure 1I). In the presence of RXP407, we observed a dose-dependent increase in MERF concentration (Figure 1J), but no change in the concentration of endogenous Met-enkephalin or Leu-enkephalin (Supplemental Figure 2A-B). RXP407 had no effect in NKO mice (Supplemental Figure 2C), ruling out non-specific effects of RXP407 on other targets. To measure the functional impact of RXP407 on synaptic transmission, we prepared acute brain slices from Drd1-tdTomato reporter mice, to guide whole-cell patch-clamp recordings from individual Drd1-expressing medium spiny neurons in the nucleus accumbens (Figure 1K and Supplemental Figure 3). Bath application of RXP407 caused long-term depression of electrically-evoked excitatory postsynaptic currents (Figure 1L), as well as an increase in the paired-pulse ratio (Figure 1K), both consistent with our prior report that elevated MERF levels reduce presynaptic glutamate release (2). Please see the Supplemental Methods for methodology description.

Our results suggest that the ACE N-terminal catalytic domain is the primary site of MERF degradation in brain tissue, and that N-terminal domain inhibition is sufficient to reduce degradation of this specific endogenous opioid peptide. This conclusion is further supported by structural modeling of the probable conformation of MERF in the ACE N-terminal catalytic domain (Supplemental Figure 4), with productive catalysis due to efficient transition state stabilization, as well as conserved active site interactions previously established for domain selectivity (6). Pharmacological inhibition of the ACE N-terminal catalytic domain thus presents a novel strategy to enhance endogenous opioid signaling in the brain. Critically, we have previously shown that central ACE inhibition does not have obvious rewarding effects, and in fact attenuates the rewarding properties of exogenous opioids like fentanyl (2). The engagement of endogenous opioid signaling without corresponding risk of misuse would be a major advance for opioid-based pharmacotherapy, with translational potential for various neuropsychiatric conditions (see Supplementary Text). Our current work will guide the development of more specific ACE inhibitors that target the primary site of MERF degradation in the brain, resulting in novel pharmacotherapies that target the endogenous opioid system with better efficacy and fewer side effects.

Animal studies were approved by the IACUC.

