## Supplemental Material for "Inhibition of the angiotensin-converting enzyme N-terminal catalytic domain prevents endogenous opioid degradation in brain tissue"

### Supplemental Text

Patients taking ACE inhibitors as a treatment for hypertension also experience unexpected positive effects, including relief from depression (1-6), improved quality of life (7, 8), and slower cognitive decline with age (9, 10). These effects appear to require central ACE inhibition in the brain, because they are not observed in patients taking ACE inhibitors that do not cross the blood-brain barrier, or patients with hypertension managed to a similar degree by other classes of medication.

### Supplemental Methods

#### *Sex as a Biological Variable*

All experiments included comparable numbers of female and male mice, with similar effects observed in both sexes.

#### *Subjects*

Experiments were performed using wild-type C57Bl/6J mice; “NKO” mice with point mutations that inactivate the N-terminal catalytic domain of ACE (11); “CKO” mice with point mutations that inactivate the C-terminal catalytic domain of ACE (12), and Drd1-tdTomato BAC transgenic reporter mice (13). Sample collection for liquid chromatography-tandem mass spectrometry occurred at 6-20 weeks of age, while whole-cell patch-clamp electrophysiology recordings occurred at 4-8 weeks old. Experimental procedures were conducted between 0900h – 1700h. Mice were housed in groups of 2-5 per cage, on a 12 hour light cycle (0600h – 1800h) at ~23ºC with food and water provided ad libitum.

#### *Drugs*

RXP407 (14) and RXPA380 (15) were synthesized by the More lab at the University of Minnesota, using previously reported methods (16). Identity and purity of the compound were confirmed by NMR and reverse phase HPLC analysis, respectively. LCMS standards were purchased from

Bachem and included Met-enkephalin-Arg-Phe (MERF), Met-enkephalin acetate salt, and Leu-enkephalin acetate salt.

*Sample Collection for Liquid Chromatography-Tandem Mass Spectrometry*

As previously described (17), individual coronal brain slices containing the nucleus accumbens and dorsal striatum (300 μm thick) were bathed in 100 μL aCSF containing (in mM): 119 NaCl, 26 $NaHCO_3$, 11 glucose, 2.5 KCl, 1 $NaH_2PO_4$-$H_2O$, 2.5 $CaCl_2$-$2H_2O$, 1.3 $MgSO_4$-$7H_2O$. For some experiments, brain slices were incubated in a saturating concentration of exogenous MERF (10-50 nM) for 20 minutes. We used Met-enkephalin production to measure degradation of exogenous MERF by ACE, since spontaneous release of endogenous Met-enkephalin is negligible under baseline conditions in the absence of stimulation (17, 18). For other experiments, brain slices were stimulated for 20 minutes using a high concentration of KCl (50 mM), to evoke the release of endogenous MERF and other peptides. To control for potential variability related to the rostral-caudal location of each coronal brain slice, drug treatments were counterbalanced to equally represent all anatomical locations in each treatment group. At the conclusion of each experiment, extracellular fluid was collected and stored at -80ºC until analysis.

*Liquid Chromatography-Tandem Mass Spectrometry (LC-MS/MS)*

Samples underwent a modified desalting protocol (19) with C18-material stage tips (SP301, Thermo Scientific), were eluted with 200 μL solvent (40:60:0.1% water:acetonitrile:trifluoroacetic acid), and dried via speed vacuum overnight. Samples were reconstituted in 12 μL solvent (98:2:0.1% water:acetonitrile:formic acid). Desalted concentrated samples underwent targeted proteomic identification and quantification based on selected reaction monitoring (SRM) and liquid chromatography-tandem mass spectrometry. Samples (5 μL) were injected onto a home-packaged analytical C18 reverse phase column (Phenomenex, Torrance, CA) and subsequently eluted with solvent A (0.1% Formic Acid [FA] in water) and solvent B (0.1% FA in ACN) with the following gradient profile: 0 - 5 min 2% solvent B flow rate at 1μL/min; 5 - 5.5 min 2% solvent B flow rate at 1 - 0.3μL/min; 5.5 - 15 min 2 - 35% solvent B flow rate at 0.3μL/min; 15 - 15.5 min 35 - 90% solvent B flow rate at 0.3 - 1μL/min; 15.5 - 19 min 90% solvent B flow rate at 1μL/min; 19 - 19.5 min 90 - 2% solvent B at flow rate 1μL/min; 19.5 - 22 min 2% solvent B at flow rate 1μL/min. Mass spectrometry detection was obtained on a TSQ Quantiva Triple Quadrupole (Thermo Scientific) in positive nanospray ionization mode. Mass spectrometry conditions were: spray voltage 2.0 kV, ion transfer tube temperature 350 °C, collision energy 4 – 29.2 V, and collision gas (argon) pressure 1 mTorr. Resolution settings were 0.7 Da (full width at half-maximum) for both quadrupoles and transition dwell times were 15 ms. Standards for absolute peptide quantification (10 pM, 50 pM, 100 pM, 500 pM, 1 nM, 5 nM, 10 nM) were injected after experimental samples and contained: Met-enkephalin, Leu-enkephalin, and MERF.

Skyline (MacCoss Lab) was used to empirically determine SRM transitions for all peptide standards and quantitative data processing. SRM transitions corresponding to the five largest integrated peaks were selected for targeted proteomic analysis, and precursor / product transitions with the largest peak area were used for absolute peptide quantification as derived from calibration curves: Met-enkephalin (574.2330 / 278.1135), Leu-enkephalin (556.2766 / 278.1135), MERF (439.2049 / 714.3392). Peaks were manually inspected to ensure correct detection and integration for each peptide per sample.

*Whole-Cell Patch-Clamp Electrophysiology*

To perform whole-cell patch-clamp recordings from D1-MSNs, Drd1-tdTomato mice were deeply anesthetized with isoflurane and perfused with 10 mL ice-cold sucrose cutting solution containing (in mM): 228 sucrose, 26 $NaHCO_3$ , 11 glucose, 2.5 KCl, 1 $NaH_2PO_4$-$H_2O$, 7 $MgSO_4$-$7H_2O$, 0.5 $CaCl_2$-$2H_2O$. Mice were subsequently decapitated and brains were quickly removed then placed in ice-cold sucrose cutting solution. Coronal slices (240 μm thick) containing nucleus accumbens (NAc) were collected using a vibratome (Leica VT1000S) and allowed to recover submerged in a holding chamber with artificial cerebral spinal fluid (aCSF) containing (in mM): 119 NaCl, 26 $NaHCO_3$, 11 glucose, 2.5 KCl, 1 $NaH_2PO_4$-$H_2O$, 2.5 $CaCl_2$-$2H_2O$, 1.3 $MgSO_4$-$7H_2O$. Slices recovered in warm aCSF (33ºC) for 10 min and then equilibrated to room temperature for at least 45 min before use. Slices were transferred to a submerged recording chamber and continuously perfused with aCSF at a rate of 2 mL/min at room temperature. All solutions were continuously oxygenated (95% $O_2$ /5% $CO_2$ ). Whole-cell voltage clamp recordings from D1-MSNs in the NAc core were obtained under visual control using IR-DIC optics from an Olympus BX51W1 microscope and distinguished by expression of tdTomato. Patched cells were voltage-clamped at -70 mV. Borosilicate glass electrodes (3-4 MΩ) were filled with (in mM): 120 $CsMeSO_4$ , 15 CsCl, 10 TEA-Cl, 8 NaCl, 10 HEPES, 5 QX-314, 4 ATP-Mg, 5 EGTA, and 0.3 GTP-Na (pH 7.2-7.3). Excitatory synaptic transmission was pharmacologically isolated using $GABA_A$ receptor antagonist picrotoxin (50 µM, Tocris). Excitatory postsynaptic currents (EPSCs) were electrically evoked locally using bipolar stimulating electrodes (ISO-flex, AMPI), with paired-pulses of stimulation delivered every 30 seconds at a 50 ms inter-pulse interval. Upon stabilization (usually 10-20 min after break-in), cells were recorded at baseline for 10 min, followed by bath application of RXP 407 (10 µM). Recordings were performed using a MultiClamp 700B (Molecular Devices), filtered at 2 kHz, and digitized at 10 kHz. Data acquisition and analysis were performed online using Axograph software. Series resistance was monitored continuously, and experiments were discarded if resistance changed by ≥20%.

**Molecular Docking**

Molecular docking studies were performed using the Maestro module of Schrödinger 2025-2 as described previously (20), to determine the potential binding conformation of MERF within the ACE active site. Briefly, the crystal structures of N- and C-terminal domains of ACE with synthetic ligands, RXP407 (PDB ID: 3NXQ) and RXPA380 (PDB ID: 2OC2), respectively, were subjected to a protein preparation workflow. Protein structure was preprocessed by adding the missing hydrogen atoms, side chain residues, and deleting water molecules beyond 5 Å from the co-crystallized ligand. A physiological protonation state was generated using PROPKA at pH 7.0, and the hydrogen bond assignments were optimized using the OPLS4 force field to achieve a low energy and stable protein structure. The receptor grid was generated within a 15 $Å^3$ box of the ligand using the Receptor Grid Generation tool. For preparation of the ligand, MERF was constructed using 3D builder and optimized by using LigPrep to generate different ionization and tautomeric conformers at pH 7 ± 2. Different generated conformers of MERF were docked in the generated receptor grids using the Glide module of Maestro with the Extra Precision (XP) algorithm. Post-docking minimization and strain correction were performed to further refine the generated poses and determine interaction profiles within the active site of each terminal domain of ACE.

*Statistical Analysis*

Data were analyzed using ANOVA models in GraphPad Prism 10, and significant main effects were further analyzed using Dunnett's post-hoc test. The Type I error rate was set to $\alpha = 0.05$ (two-tailed) for all comparisons. In the figures, significant ANOVA effects and post-hoc tests are denoted by black asterisks that indicate *p<0.05, **p<0.01, ***p<0.005, and ****p<0.0001. All data are presented mean ± SEM, with open and closed circles indicating data points from female and male mice, respectively.

**Study Approval**

All procedures were approved by the Institutional Animal Care and Use Committee at the University of Minnesota, and conformed to the National Institutes of Health Guidelines for the Care and Use of Laboratory Animals.

### Data Availability

Raw data for all figures can be found in the Supporting Data Values file.


### Acknowledgments

This work was supported by the University of Minnesota's MnDRIVE (Minnesota's Discovery, Research, and Innovation Economy) initiative (F.H., P.E.R.); an Interdisciplinary Doctoral Fellowship from the University of Minnesota Graduate School (F.H.); a Faculty Research Development Grant from the University of Minnesota Office of Academic Clinical Affairs (S.S.M., P.E.R.); and NIH grants DA060664 (J.L.S.), DA056331 (S.S.M., P.E.R.), and DA056675 (S.S.M., P.E.R.). Mass spectrometry was carried out in the Analytical Biochemistry Shared Resource of the Masonic Cancer Center, supported in part by the U.S. National Institutes of Health and National Cancer Institute (Cancer Center Support Grant CA-77598). Schematics were created with BioRender.com.

### Supplemental Figures

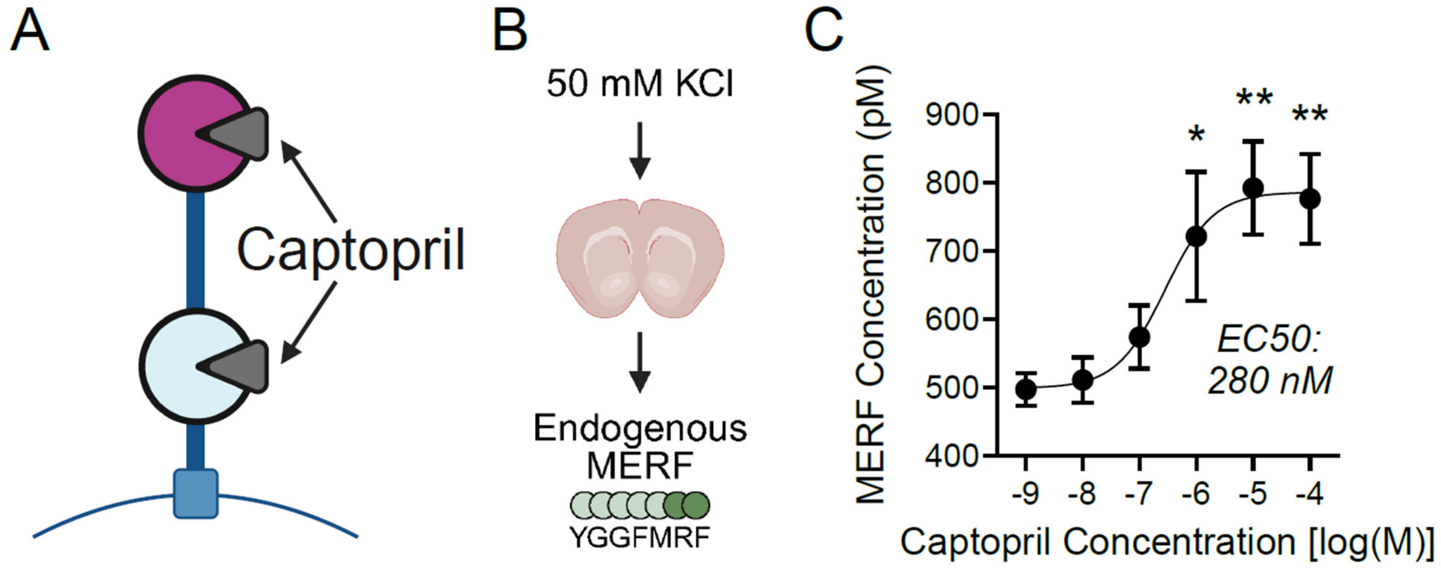


**Supplemental Figure 1. Pharmacological inhibition of angiotensin-converting enzyme (ACE) with captopril prevents MERF degradation.** (**A**) Schematic showing pharmacological inhibition of both catalytic domains of ACE by captopril, a prototypical ACE inhibitor with little domain selectivity. (**B**) Experimental design: stimulation of brain slices with 50 mM KCl to release endogenous MERF. (**C**) Effects of captopril on extracellular concentration of MERF in wild-type brain slices (n=9-12/dose). ANOVA main effect of Dose: $F_{5,56}$ = 5.38, p = 0.0004; *p<0.05, **p<0.005, Dunnett's post-hoc test versus lowest captopril concentration. Graph displays mean ± SEM.

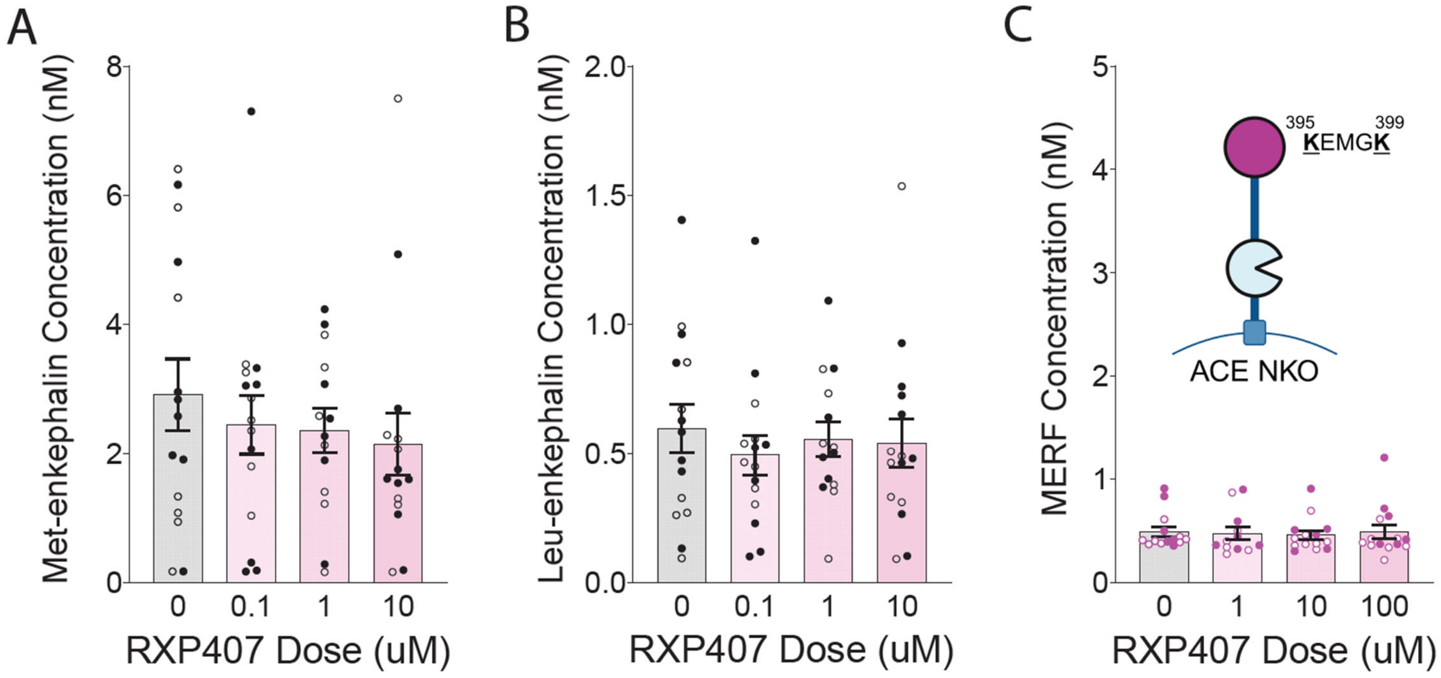


**Supplemental Figure 2. Control experiments demonstrating the specificity of pharmacological inhibition of the angiotensin-converting enzyme (ACE) N-terminal catalytic domain with RXP407.** (**A-B**) No effect of RXP407 on extracellular concentration of Met-enkephalin (**A**) or Leu-enkephalin (**B**) in brain slices from WT mice (n=15-16 per dose). (**C**) No effect of RXP407 on extracellular MERF concentration in brain slices (n=15-16 per dose) from "NKO" mice carrying point mutations that inactivate the N-terminal catalytic domain of ACE. Graphs display mean ± SEM, with individual data points from females and males shown as open and closed circles, respectively.

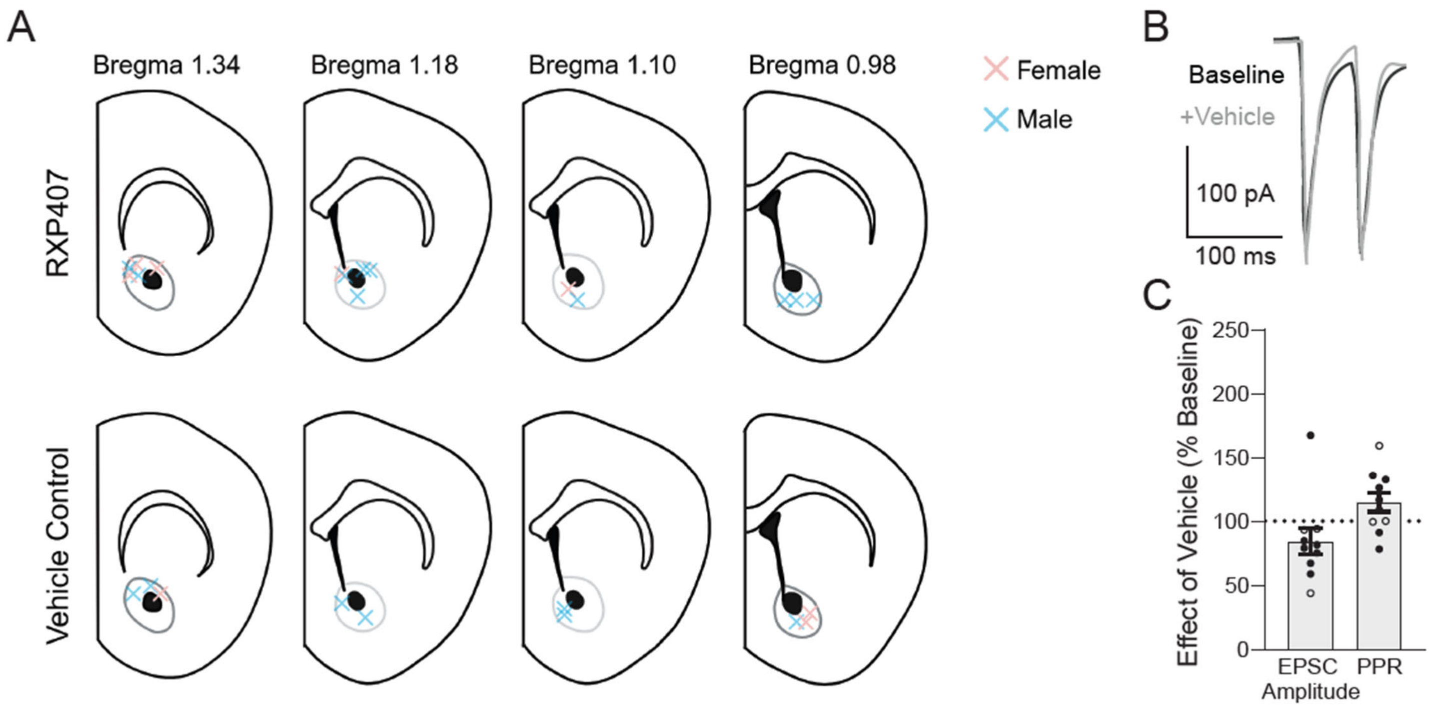


**Supplemental Figure 3. Additional information related to whole-cell patch-clamp recordings for electrophysiology experiments.** (**A**) Location of cells recorded in the nucleus accumbens core subregion, from female mice (pink X) and male mice (blue X) and treated with RXP407 (top row) or vehicle (bottom row). (**B**) Example traces showing electrically-evoked excitatory postsynaptic currents (EPSCs) before and after treatment with vehicle. (**C**) No significant effects of vehicle on EPSC amplitude or paired-pulse ratio. Graphs display mean ± SEM, with individual data points from females and males shown as open and closed circles, respectively.

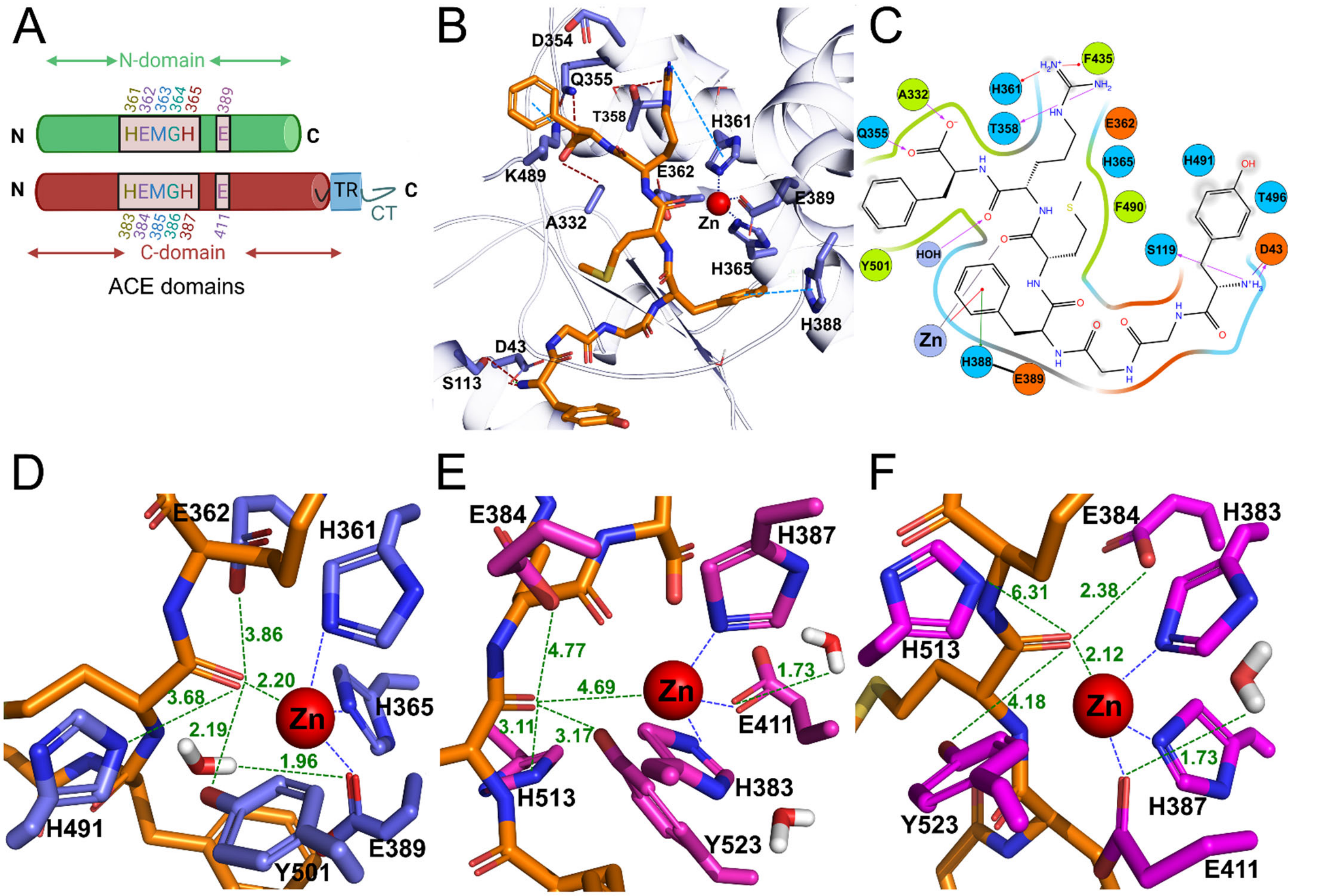


**Supplemental Figure 4. Molecular interactions of Met-enkephalin-Arg-Phe (MERF) with active site residues of N- and C-terminal domains of angiotensin-converting enzyme (ACE).** (**A**) Alignment of key metal binding residues within N- and C-terminal ACE domains. TR: transmembrane region, CT: C-terminal tail. (**B**) Docked conformation of MERF showing occupancy and key hydrogen bonds within the catalytic channel of the N-terminal domain. (**C**) 2D diagram displaying active site interactions of MERF with $Zn^{2+}$ and the surrounding residues of the N-terminal domain. (**D**) Magnified view of MERF interactions with catalytic residues within the N-terminal domain. The enzyme-substrate complex in the N-terminal domain displays chelation of the scissile amide carbonyl (between $Met^5$ and $Arg^6$) by $Zn^{2+}$, promoting attack by the active site water bound to Glu389 present in the proximity. The resultant oxyanion tetrahedral intermediate is stabilized by hydrogen bond interactions with His491 and Tyr501 (21). (**E-F**) Two predicted binding conformations of MERF with the C-terminal domain catalytic residues. The lowest energy conformation is induced by the S2' hydrophobic pocket, reorienting MERF in a conformation less favorable for catalysis. The second conformation displayed increased distances from the catalytic residues, potentially reducing the enzymatic efficiency.
